# ‘Silent’ mitral cells emerge from common feature subtraction

**DOI:** 10.64898/2026.10.22.761107

**Authors:** Sina Tootoonian, Mihaly Kollo, Andreas T. Schaefer

## Abstract

Whole-cell recordings from olfactory bulb mitral/tufted cells in awake mice revealed an inverse relationship between baseline activation and odour responses wherein cells that were spontaneously active tended to be inhibited by odours, while ‘silent’ cells, quiet at baseline, tended to have vigorous odour responses. In this work we provide a normative explanation for these findings by extending previous sparse coding models of the olfactory system to include ‘common feature subtraction,’ an odour-triggered variant of background subtraction in which the receptor contribution of features common to the odours being discriminated is subtracted. Using a circuit-level formulation of this computation consistent with the anatomy of the olfactory bulb we show how baseline activity cancels the effect of the common olfactory features on receptor activations, promoting inference of the target odours. We provide simulation results showing qualitative agreement with experimental findings. The model also predicts in closed form, without fitting, the slopes of the relationships between a cell’s baseline and its common input and between its baseline and its odour response, and these predictions agree with simulations. Our model makes the experimentally testable prediction that a cell’s baseline activation is not an intrinsic property, but reflects the *negative* of its recent activity.

## 1 Introduction

Modern neural recording techniques are often biased towards neurons that are highly active during the baseline state. To avoid such biases Kollo et al. [KSA^+^14] used blind whole-cell recordings to study the responses of mitral/tufted cells in the olfactory bulb of awake mice during passive odour exposure or an odour discrimination task. Interestingly, they found that most of the odour-evoked spiking activity was due to ‘silent’ cells, so named because of their dormancy during baseline periods. Conversely, cells that were active during the baseline period tended to be inhibited by odours. How do such responses emerge, are they due to intrinsic properties or are they learned, and what, if any, computational role do they play in olfactory processing?

In this work we propose answers to these questions by showing how both types of responses emerge as a result of olfactory *common feature subtraction*, whereby receptor activations common to recent odours are subtracted to facilitate discrimination. We begin by briefly reviewing the olfaction-as-inference framework upon which we build our work. We then show how it can be extended to include feature subtraction, and how doing so implies an inverse relationship between background and odour-evoked activity. We verify this relationship with simulation results, which show good qualitative agreement with those of [KSA^+^14]. Finally, we suggest simple changes to existing experimental protocols that could test the predictions of our model.

## 2 Results

### 2.1 Background: Olfaction as inference

Here we briefly summarize the relevant components of the olfaction-as-inference framework of [TSL22] on which our work is based; see [KR11, GBB^+^17, ZVMT^+^23] for related formulations. Our new contribution begins in Sec. 2.2.

Odours are modeled as high-dimensional molecular concentration vectors **x** = {*x*_1_, *x*_2_, …, *x*_*N*_ }, where component *x*_*i*_ encodes the concentration of the *i*’th molecular species. Components are assumed to come from the distribution

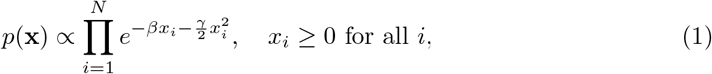

where *β* encourages sparsity and *γ* penalizes high concentrations.

We assume that a glomerulus *y*_*i*_ is excited linearly by every component *x*_*j*_ of an odour **x** according to glomerulus-specific affinities *A*_*ij*_, so that the total excitation produced by all components in the odour is

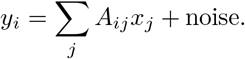

We take the noise to be normally distributed with variance *σ*^2^, yielding the likelihood

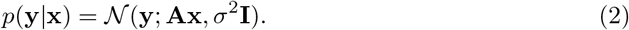

This likelihood can be combined with the prior in Eq. (1) to form a posterior. We assume that the olfactory system performs inference by maximizing this posterior given receptor inputs. This procedure is known as maximum *a posteriori* (MAP) inference, and reports

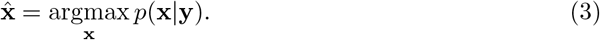

This computation can be implemented as the fixed-point of the dynamics

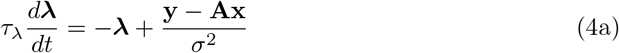

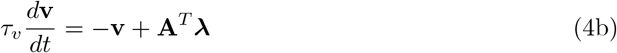

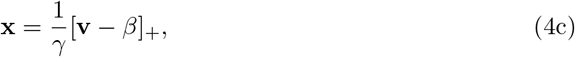

where [*x*]_+_ ≜ max(0, *x*) indicates linear rectification. The dimensionality and connectivity implied by these dynamics makes it natural to map ***λ*** to mitral/tufted cells and **x** to granule cells. Because ***λ*** is not rectified and can take either sign, we interpret it as a membrane-potential-like quantity rather than a firing rate; where we compare to spike counts (Figure 8) we convert it explicitly. See [TSL22] for details.

### 2.2 Common feature subtraction

To motivate our present work, consider discriminating a sequence of wines by their odours alone. This is shown schematically in Fig. 1A. The odour profiles will have much in common simply by virtue of their being emitted by wines. In addition to these *common features*, the smell of each wine will also contain *unique features* that distinguish each wine from the others. It would clearly be useful for the discriminator to be able to subtract the olfactory features common to all the wines to better focus on the unique features that determine the discrimination. Mice can indeed detect target odours embedded in variable backgrounds [RHKM14]. Note importantly that this is a distinct computation from subtracting whatever background odour is present in the room, which can likely be done through receptor adaptation alone. The computation of interest here is subtracting the olfactory features present in the wines from a sequence of sniffs, some of which may contain only background air. In this section we will describe how this can be done through an odour-triggered variant of background subtraction that we call *common feature subtraction* (CFS).

**Fig 1.**
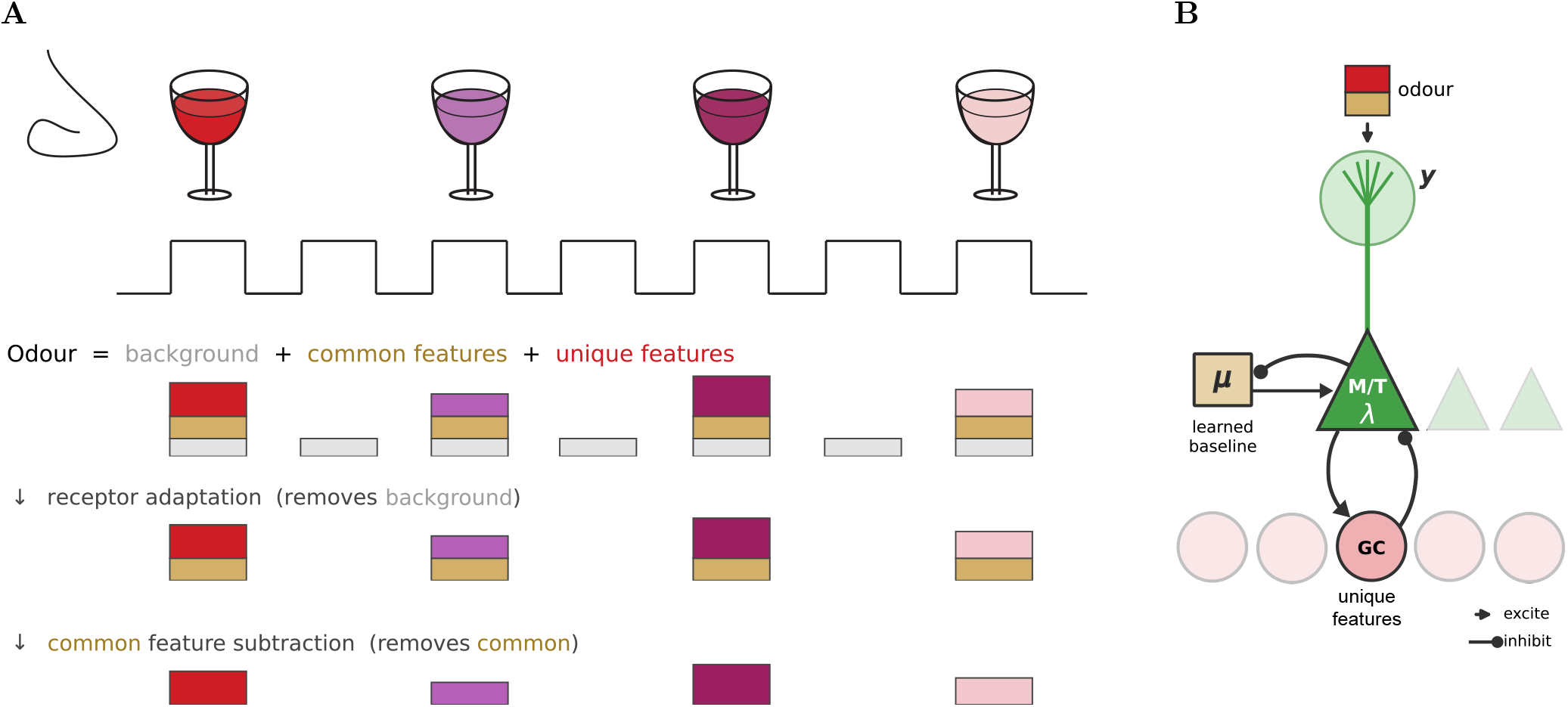
Common feature subtraction. **(A)** A discriminator samples a sequence of odours in sniffs (top; four wines, colours). Each odour is the sum of an ever-present *background* (air, gray), a *common feature* shared by the odours (tan), and an odour-specific *unique feature* (colour) (second row). Receptor adaptation removes the constant background (third row); common feature subtraction removes the common feature, leaving only the unique features that distinguish the odours (bottom row). **(B)** Circuit. The odour excites the glomerular tuft (receptor input *y*) of a mitral/tufted (M/T) cell. Granule cells (GC) for the unique features form reciprocal excitatory/inhibitory synapses with the M/T cell, and a learned baseline *µ*, added at the M/T cell, cancels the common feature.

To capture scenarios like the wine discrimination we make the following modifications to the olfaction-as-inference framework summarized in Sec. 2.1:

1. We add a temporal component to odour presentation and split time into consecutive *bouts* approximately one second long, corresponding roughly to individual sampling episodes (a sniff or a short burst of sniffing).
2. The odour in each bout *n* is the sum of common features **c** and unique features **x**,

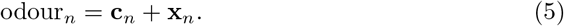
3. Common features are independent of unique features, so their joint probability over bouts factorizes as

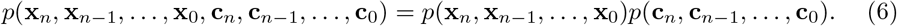
4. Common features change slowly and depend on the previous bout,

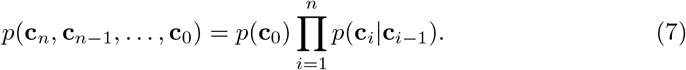
5. Unique features change quickly and are generated independently every bout,

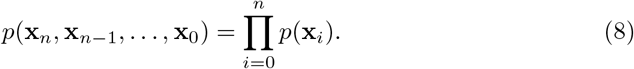
6. We allow for the possibility that a bout contains only air instead of odour, indicated by the binary variable *o*_*n*_. We treat this variable as given. Natural odour plumes are highly intermittent, consisting of brief, concentrated filaments separated by clean air [PBAP14], so *o*_*n*_ reflects a physical fact about the arrival of an odour packet rather than an inference about odour identity, and can be computed cheaply, e.g. by thresholding the summed activation of the hundreds of available glomeruli (Sec. 4.6); in the simulations reported here *o*(*t*) is supplied directly. The animal observes odours indirectly via their receptor activations **y**_*n*_. As in [TSL22], we assume that odours excite receptors linearly. Our next assumption is that
7. Receptor adaptation has already removed the excitation by any odours present in the air, so the resulting excitation only contributes a noise term. Therefore, in bout *n*, the receptor excitations are

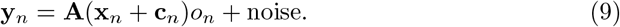

Thus, when the bout contains only air, and not also odour, *o*_*n*_ = 0 and receptor observations revert to background noise. Gating by this variable is what allows for *odour-triggered* feature subtraction (see below). Just as in [TSL22], we take the task of the olfactory system to be maximum *a posteriori* (MAP) inference of the odour that caused the receptor activations. However, our final assumption is that
8. The animal only cares about the unique odour features **x**_*n*_. Therefore, the inference task is to compute

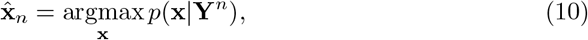

where **Y**^*n*^ denotes the full history of receptor observations

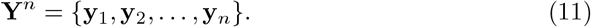

In order to infer the unique features from the receptor activations, the inference process must also account for the effect of the common features, and it is this accounting that leads to the emergence of ‘silent’ cells, as we show below.

### 2.3 Inference when odour is present

If the current bout does not contain odour then receptor activations do not provide any information about what the unique features could be. In that case, the posterior on the unique features reverts to the prior, whose peak, due to the sparsity assumption, is at **x** = **0**, so that MAP inference is computed trivially. For this reason we will focus on the more interesting case where odour is assumed present i.e. *o*_*n*_ = 1.

By Bayes’ rule, the posterior probability of the unique features **x**_*n*_ in the current bout, given that odour is present, and the history of receptor observations up to the present bout, satisfies

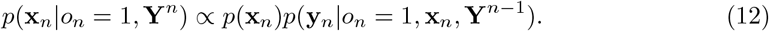

The latter term relates the most recent observation **y**_*n*_ to the previous history of observations

**Y**^*n−*1^. The current observation **y**_*n*_ depends on the common features **c**_*n*_ in the current bout, which in turn depends on the common features **c**_*n−*1_ in the previous bout. Neither of these are known, so we have to account for all possible values they could take by averaging over them, giving the double integral

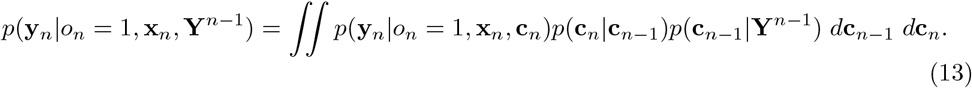

Computing this integral is complex for two reasons. The first is that **c**_*n*_ and **c**_*n−*1_ are high dimensional vectors, for which a single integral can often be intractable, not to mention a double integral. Secondly the integral requires knowing the transition model *p*(**c**_*n*_ |**c**_*n−*1_), a doubly-high dimensional object describing transitions between sparse vectors that may also be intractable to learn and use.

To avoid these problems, we recall our assumption that the animal only cares about the *unique* features **x**_*n*_. The common features **c**_*n*_ are a nuisance parameter that must be accounted for, but the precise molecular profile of **c**_*n*_ is not important to the animal. Hence instead of accounting for the common features directly in high-dimensional latent space, they can instead be accounted for through their effects on low-dimensional receptor space. From the observation model in Eq. (9), this effect is **Ac**_*n*_. We will instead model the negative of this quantity,

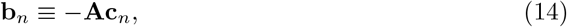

so that its mean will determine the mitral cell baseline activity; see Eq. (19) below.

The vector **b**_*n*_ is a random low-dimensional projection of a high-dimensional object and is therefore likely to have approximately Gaussian dynamics. We therefore assume that the probability distribution of receptor activations due to common features, given the history of receptor observations is Gaussian with mean ***µ***_*n*_ and variance 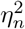,

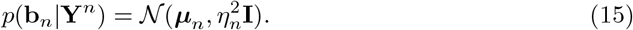

The receptor effects change slowly from one bout to the next, and we assume that these changes are also Gaussian, so

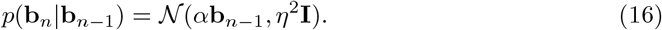

The parameter *α* determines the history dependence. When *α* = 0 the mean of receptor activations due to the common features in this bout has no relation to that in the previous bout i.e. there is no history dependence. When *α* = 1, the mean in the current bout is given by the activity in the previous bout i.e. full dependence. We assume that 0 *< α <* 1. Combining this Gaussian model of the common-feature effects with the receptor likelihood, and updating it after each bout, yields a modified mitral cell inference and an update rule for the baseline ***µ***, which we derive in Methods (Sec. 4.1).

Collecting the mitral cell dynamics in Eq. (19), and converting the discrete-time common feature update of Eq. (31) into continuous time (see Methods) yields the full system dynamics for common feature subtraction. Since the M/TC baseline is *α****µ***, with some abuse of notation we’ve redefined ***µ*** as *α****µ***. Collecting constants for clarity into

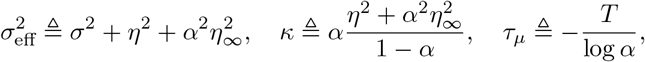

the system dynamics become

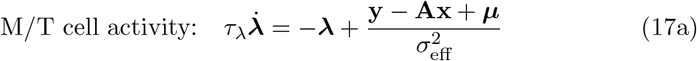

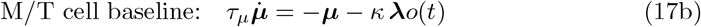

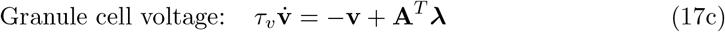

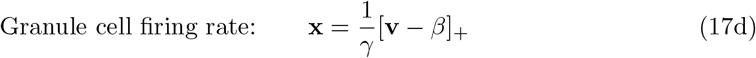

Here *T* is the average bout duration, the time-varying binary signal *o*(*t*) indicates whether odour is present, and bout-dependent variances 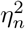 have been replaced with their steady-state values 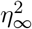 from Eq. (33).

The innovation of the present work is the introduction of the M/T cell baseline ***µ***, and the derivation of its dynamics as Eq. (17b). What makes ***µ*** the M/T cell *baseline* is that, by Eq. (17a), M/T cell activity in the absence of glomerular input **y** and granule cell input **x** converges to a value proportional to ***µ***. Note that Eq. (17a) applies the baseline at all times, not only when odour is present. This is deliberate: because *o*(*t*) is not known in advance, the cancellation of the common input must already be in place when an odour packet arrives, so the baseline acts as a standing prediction of the common input. During air the baseline’s standing drive to the granule cells, 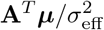, remained below the granule cell threshold *β* in the simulations reported here. What makes the baseline update *odour-triggered* is the multiplicative interaction between the M/T cell activity and the odour presence signal *o*(*t*) in Eq. (17b). Crucially, the dynamics in Eq. (17b) imply that the baseline then reflects the *negative* of the average odour-triggered activity of each M/T cell over the time window *τ*_*µ*_ = −*T/* log *α*, which we assume is long relative to the average inference time. Thus the baseline is *inversely* related to the recent odour-triggered activity of each M/T cell, and it is this property of the model which may explain several of the key experimental findings of [KSA^+^14].

### 2.4 Simulations

To demonstrate the performance of the model and to compare it to experimental observations, we next performed a series of simulations.

In Figure 2 we show the responses of granule cells and a few example mitral cells (we henceforth refer to M/T cells collectively as mitral cells for simplicity) to four odours, before (early) and after (late) common feature subtraction has been learned. Each odour is a binary mixture of a common feature and a unique feature. Early on, before the mitral cell baselines have adapted, the baselines are at zero and granule cells decode both the common feature and the unique feature. Once the system has learned the common feature, its learned baseline cancels the common drive: the granule cell for the common feature falls silent, and mitral cell baselines spread apart—some becoming persistently silent—so that only the unique feature is decoded.

**Fig 2.**
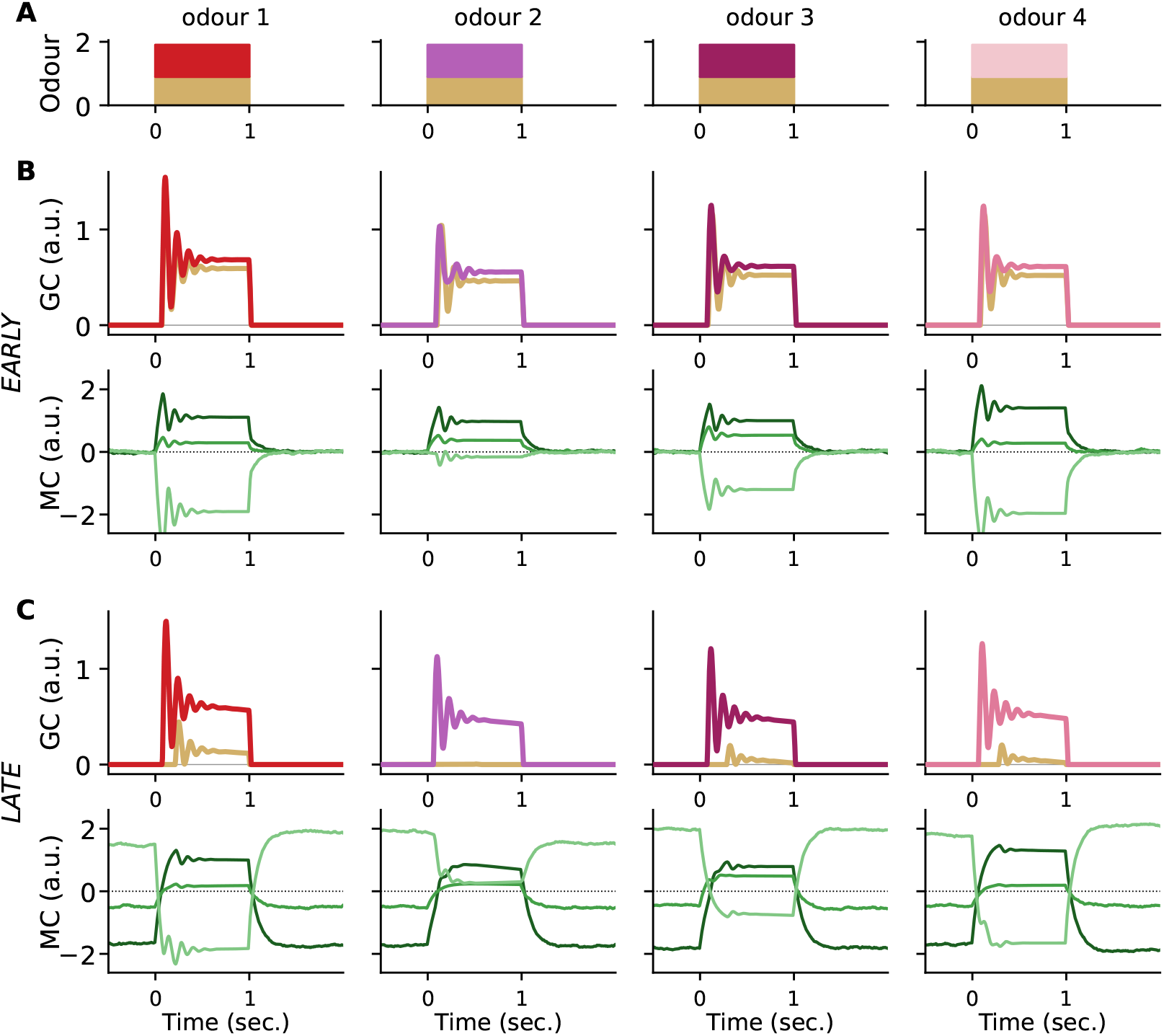
Responses before and after common feature subtraction. The same four odours (columns) before (B, early) and after (C, late) the common feature has been learned. **(A)** Each odour is a common feature (tan) plus a unique feature (colour). **(B)** Early: the granule cells (GC) decode both the common (tan) and unique (colour) features, and the three example mitral cells (MC, green) respond to the odour from baselines at zero; the baselines are zero because the baseline update has not yet been switched on (Sec. 4.3). **(C)** Late: the learned baseline has cancelled the common feature, so the common granule cell is almost silent, leaving only the unique feature, and the mitral cell baselines have spread apart, rendering some cells persistently silent. The damped oscillation at odour onset is a property of the network dynamics (time constants *τ*_*λ*_, *τ*_*v*_ and loop gain 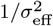) and makes no claim about sniff- or theta-frequency rhythms.

Figure 3 shows how this comes about. We follow one of the mitral cells of Figure 2—the one that ends up silent—through four presentations of the same odour, from the first (early) to one after its baseline has converged (late), with two intermediate presentations between. Once learning is switched on after the priming presentations, every odour presentation drives the cell—all odours share the common feature—and by Eq. (17b) each such activation lowers its baseline; the four columns sample this process at successive presentations of one odour. As the baseline falls, the common drive is progressively cancelled: the granule cell for the common feature responds less on each presentation and is eventually silent, while the granule cell for the unique feature is unaffected. Note that the activity of the mitral cell *during* the odour changes comparatively little over learning (by about 20% here, against a baseline shift six times larger); what changes is its baseline, and with it the division of labour between the granule cell for the common feature and the baseline in accounting for the common drive. The mitral cell itself settles into the ‘silent’ profile of [KSA^+^14]: quiet at rest, with a large odour-evoked depolarization from that baseline. The number of presentations required is set by *τ*_*µ*_ (Table 1).

**Table 1.**
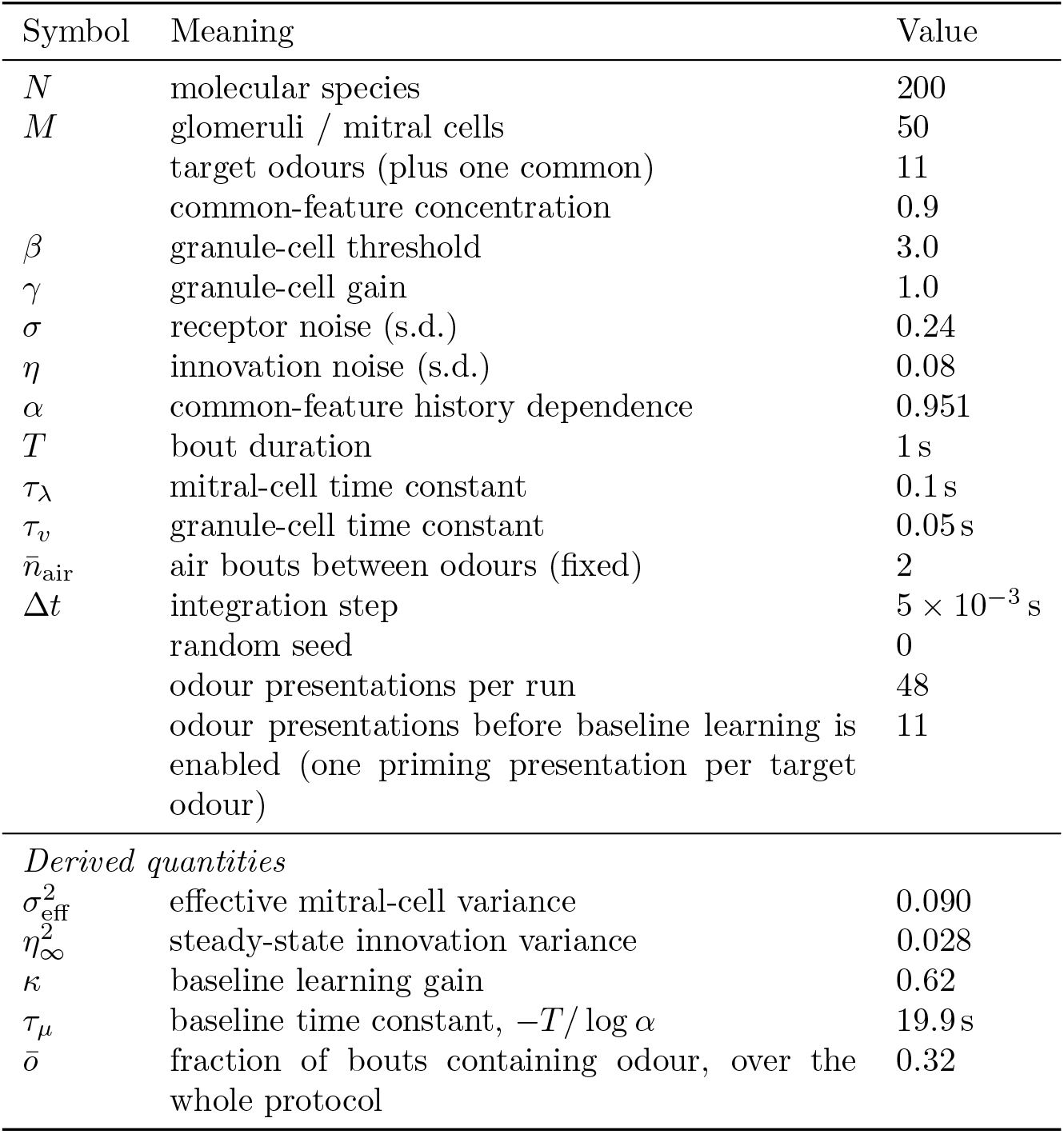
Simulation parameters for the default run (Figures 4–7). Quantities below the rule are derived from those above.

**Fig 3.**
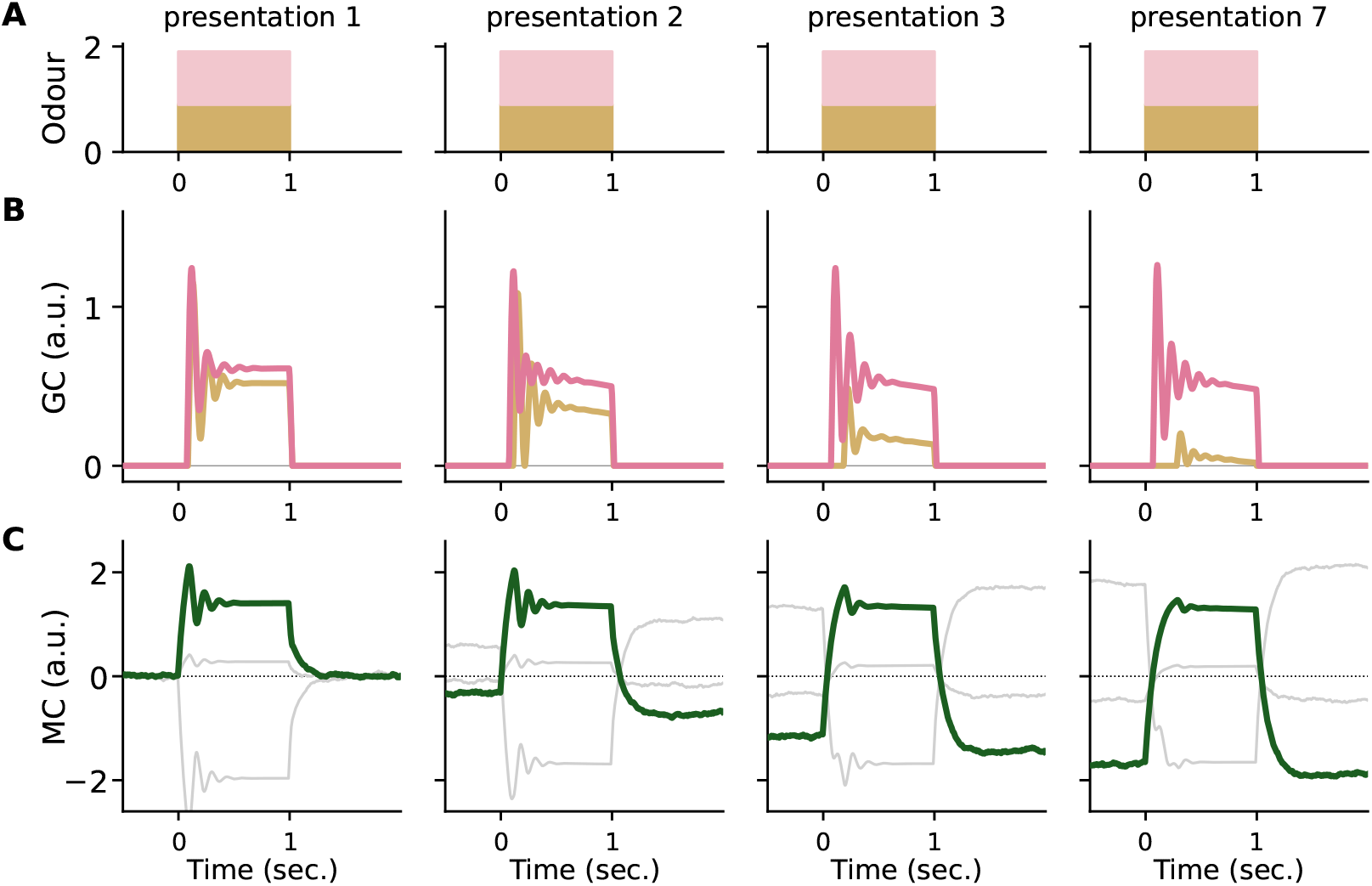
Time course of learning. One mitral cell from Figure 2 (dark green; the other two cells of that figure in grey) and the granule cells, responding to the same odour at four presentations: the first (early), two intermediate presentations, and one after the baseline has converged (late). The early and late columns are the corresponding panels of Fig. 2B and Fig. 2C; column labels give the presentation number of this odour; the other odours, which share the common feature, are presented in between. **(A)** Odour: the common (tan) and unique (colour) features. **(B)** Granule cells (GC): with each presentation the baseline cancels more of the common drive, so the common-feature granule cell responds less, until it is almost silent; the unique-feature granule cell is unaffected. **(C)** Mitral cells (MC): the baseline of the followed cell falls below zero with successive presentations, rendering it silent at rest; its activity during the odour changes comparatively little, so its odour-evoked depolarization from that baseline grows as the baseline falls.

According to Eq. (17b) the mitral cell baseline should be inversely correlated to the glomerular excitation caused by the common odour. In Fig. 4A we verify this by plotting the glomerular excitation due to the common odour and the baseline activity of each cell in response to the final 10 odour presentations, and indeed observe a pronounced negative correlation. Beyond the sign of this relationship, our derivation predicts its *slope* in closed form (Eq. (44)). To test this prediction, in Fig. 4B–F we repeat the simulation while changing, one at a time, parameters that our expression predicts should alter the slope. In each case the slope predicted from the model parameters alone, with no fitting (red line), tracks the simulated data; fitted slopes are shown alongside the predictions in each panel. Fig. 4D also shows an ungated control, discussed below.

**Fig 4.**
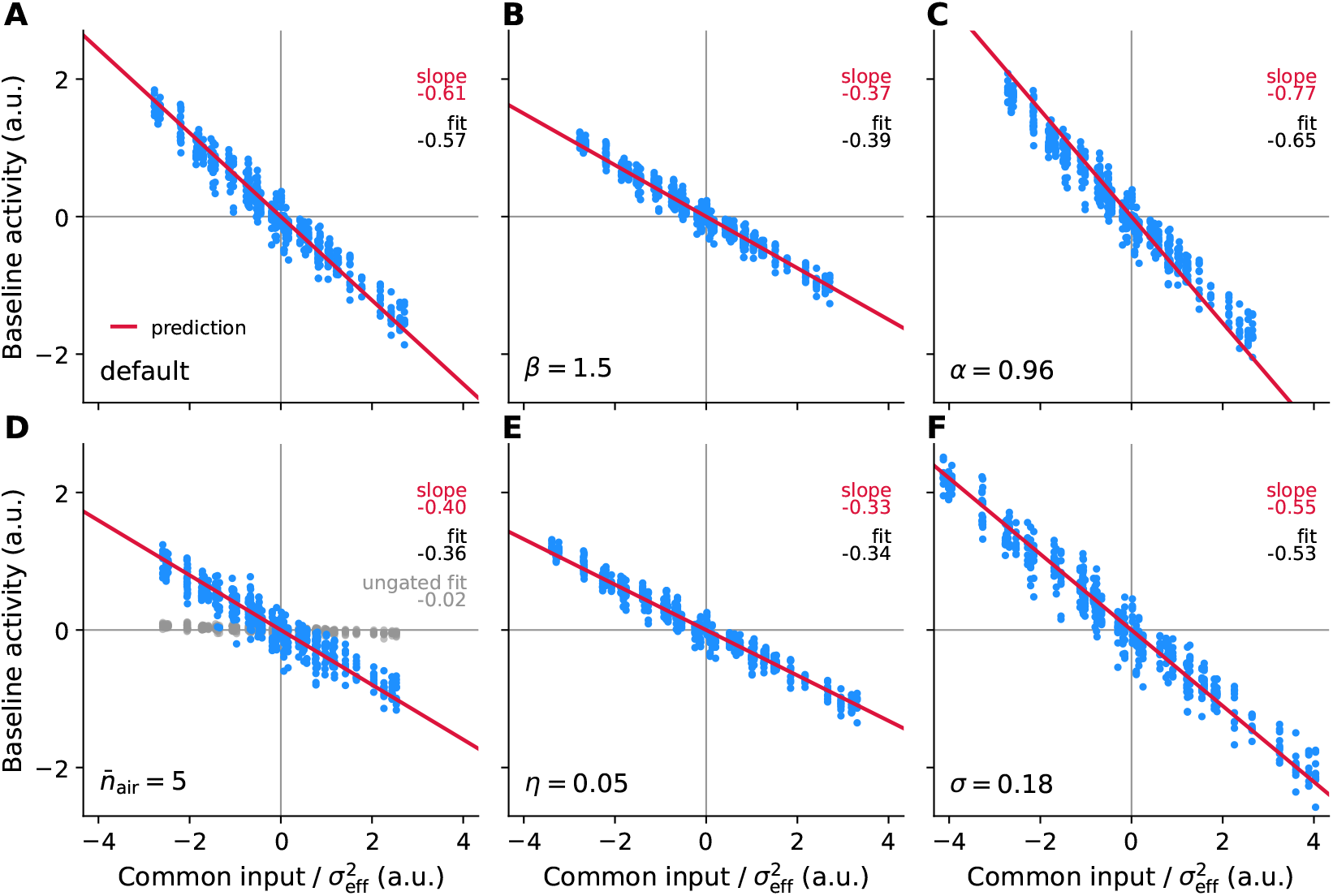
The predicted slope of the mitral-cell baseline against the common input. Each panel plots the common-odour input to a glomerulus, divided by 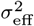 (x-axis), against the pre-odour baseline activity of the mitral cell sampling that glomerulus (y-axis), over the final 10 odour presentations; all panels share the same axes. The red line is the slope predicted by Eq. (44), computed from the model parameters with no fitting. **(A)** Default parameters. **(B–F)** As in A but with a single parameter changed from its default value (*β* = 3, *α* = 0.951, 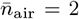, *η* = 0.08, *σ* = 0.24): **(B)** sparsity/threshold parameter *β* = 1.5; **(C)** common-feature history dependence *α* = 0.96; **(D)** inter-odour air gap 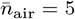; grey points show the same simulation with the baseline update applied at all times rather than only during odour (ungated; see Discussion); **(E)** common-feature innovation noise *η* = 0.05; **(F)** receptor noise *σ* = 0.18. The predicted slope tracks the simulated data across all manipulations.

Next, we systematically compared the simulated responses to those reported in [KSA^+^14]. To best allow comparison to the results reported there, we used the responses of all 50 mitral cells to the final 10 odour presentations, yielding 500 cell-odour pairs. Odour responses were defined as the mean voltage during each 1-second odour presentation, and baseline responses were the mean voltage in the 500 ms preceding odour presentation.

In Figure 5 we compared the distribution of odour-evoked responses to those in Fig. 2b of [KSA^+^14]. In that work the authors found a range of response amplitudes, with positive responses tending to be accompanied by low baseline activity and vice versa. Because the simulated responses are noiseless (Sec. 4.3), the significance-based classification of responses used in [KSA^+^14] does not apply. Instead, in Fig. 5C we plotted the odour-evoked responses against the baseline responses, revealing a clear negative correlation, in qualitative agreement with the results in [KSA^+^14]. As with the baseline, our derivation predicts the slope of this relationship in closed form (Eq. (45); Methods). The red line in Fig. 5C is this prediction, computed from the model parameters with no fitting; it passes through the per-cell mean responses, with individual cell-odour pairs scattered about it.

**Fig 5.**
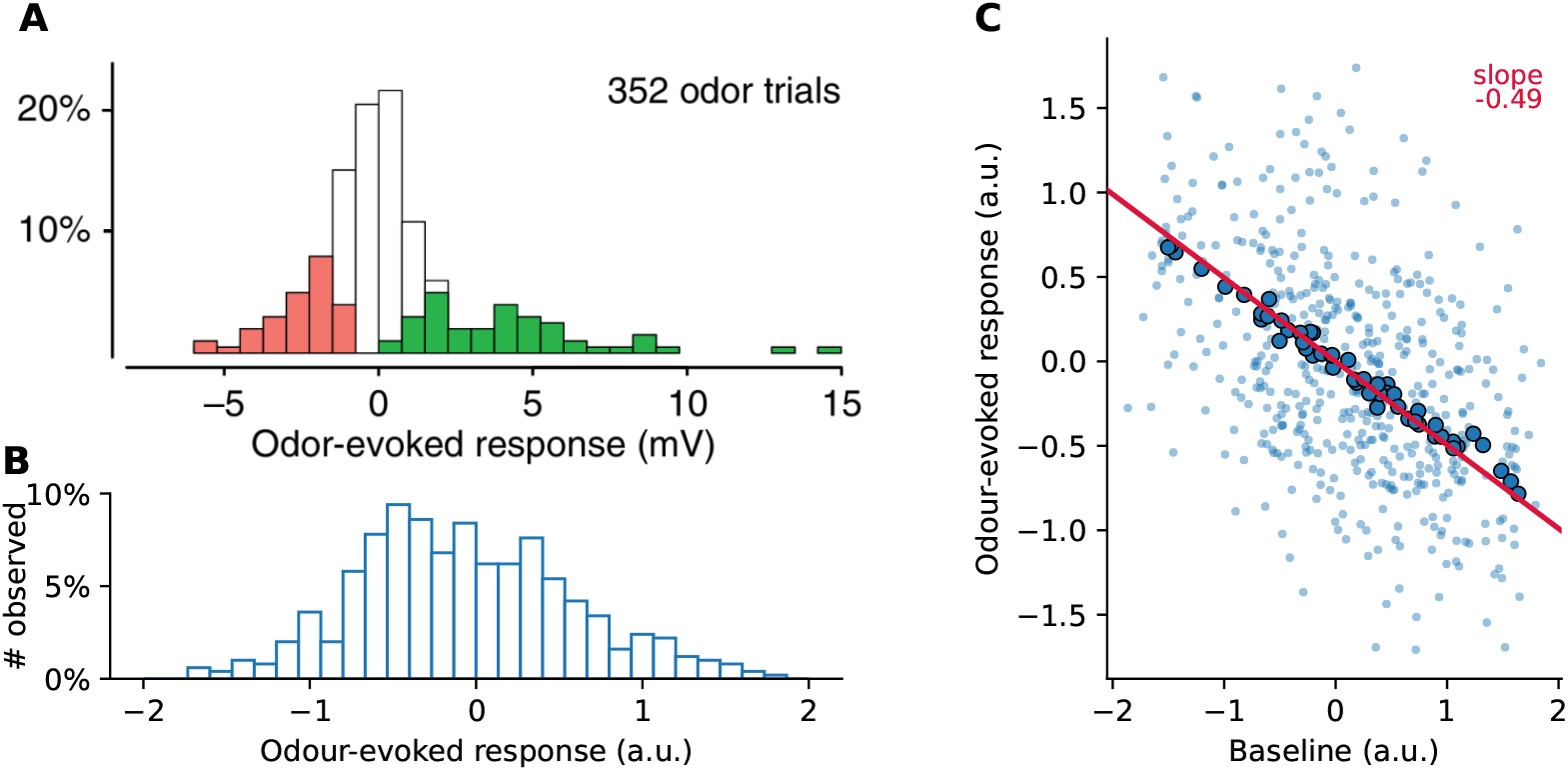
Response distributions. **(A)** Experimentally observed distribution of odour-evoked responses. Green and red bars indicate significant depolarizing and hyperpolarizing responses, respectively. Adapted from Fig. 2b of [KSA^+^14]. **(B)** Distribution of the mean voltage during the odour presentation period computed over 500 odour-cell pairs. **(C)** Relation of the odour-evoked responses in panel B (y-axis) to the mean baseline activity in the 500 ms before odour onset (x-axis). Baseline activity was inversely related to odour-evoked responses. Outlined points: per-cell means over the 10 presentations. Red line: slope predicted by Eq. (45) from the model parameters, with no fitting; it passes through the per-cell means (slope of the per-cell means *−*0.45, predicted *−*0.49).

In Fig. 2c of [KSA^+^14] the authors compare the distribution of odour responses grouped by the pre-odour firing rate. They find that higher pre-odour firing rates result in lower odour responses. In Fig. 6B we performed a similar analysis, grouping the responses by their relation to the median baseline response and comparing the odour-evoked response distributions of the resulting two groups. In agreement with [KSA^+^14], we find that higher pre-odour voltages result in lower odour-evoked responses.

**Fig 6.**
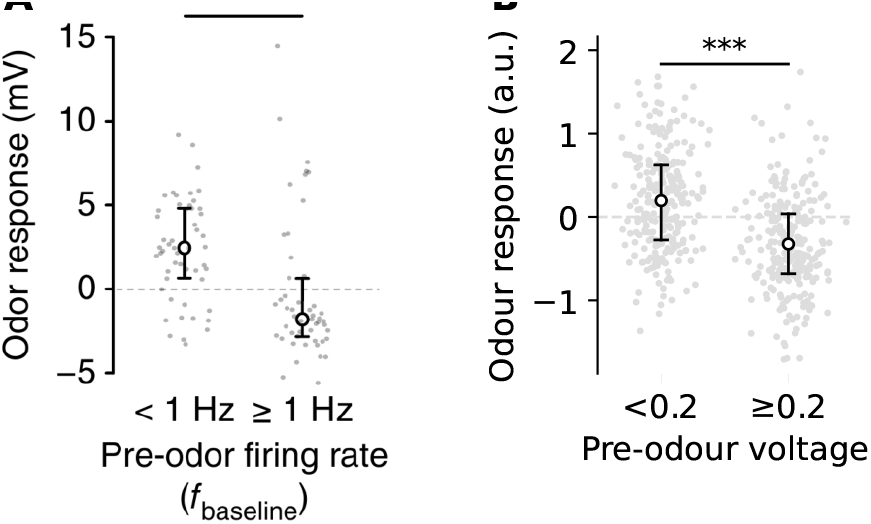
Odour responses grouped by pre-odour activity. **(A)** Experimentally observed distributions of odour responses, grouped by pre-odour firing rate. Circles are means, whiskers are 25th and 75th percentiles. Higher pre-odour firing rates meant lower odour responses (*P <* 0.001, Mann-Whitney U test). Adapted from Fig. 2c of [KSA^+^14]. **(B)** Distribution of simulated odour responses in 500 cell-odour pairs, grouped by mean activity in 500 ms prior to odour onset. Circles are means, whiskers are 25th and 75th percentiles. Higher pre-odour activity meant lower odour responses. To avoid treating the 10 presentations of each cell as independent, statistics were computed on per-cell means: cells split at the median per-cell baseline (*n* = 25 per group) differed in mean odour response (*P <* 10^*−*8^, Mann-Whitney U-test), and per-cell mean baseline and mean response were strongly anticorrelated (Spearman *ρ* = *−*0.99, *n* = 50 cells).

Next, in Figure 2d of [KSA^+^14] the authors show how the baseline voltage relates to the distribution of odour-evoked responses by plotting the cumulative distributions of odourevoked responses for a range of baseline voltages. In Fig. 7B we perform the same analysis for our simulated responses, showing that higher baseline voltages lead to lower odour-evoked responses, in qualitative agreement with the results in [KSA^+^14].

**Fig 7.**
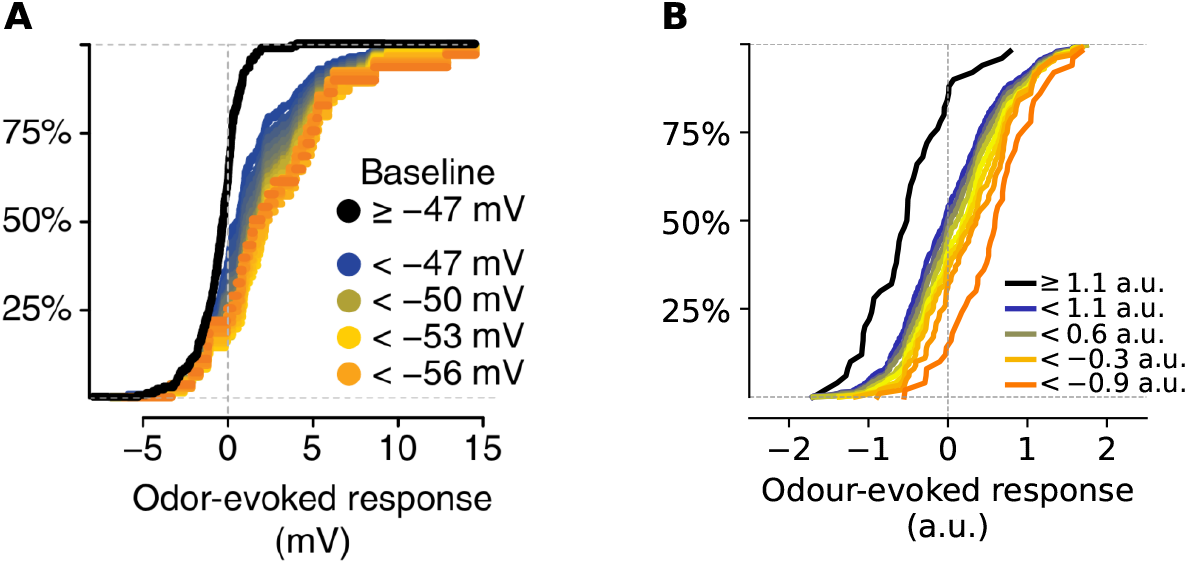
Cumulative distribution of odour-evoked responses. **(A)** CDF of experimentally observed odour-evoked responses, grouped by baseline voltage. Higher baseline voltages result in lower odour-evoked responses. Adapted from Fig. 2d of [KSA^+^14]. **(B)** CDF of simulated odour-evoked responses, grouped by baseline in the same way as A: the black curve contains the cell-odour pairs with baseline at or above the 90th percentile, and each coloured curve contains all pairs with baseline below one decile threshold (only alternate thresholds are labelled). Higher baselines result in lower odour-evoked responses.

**Fig 8.**
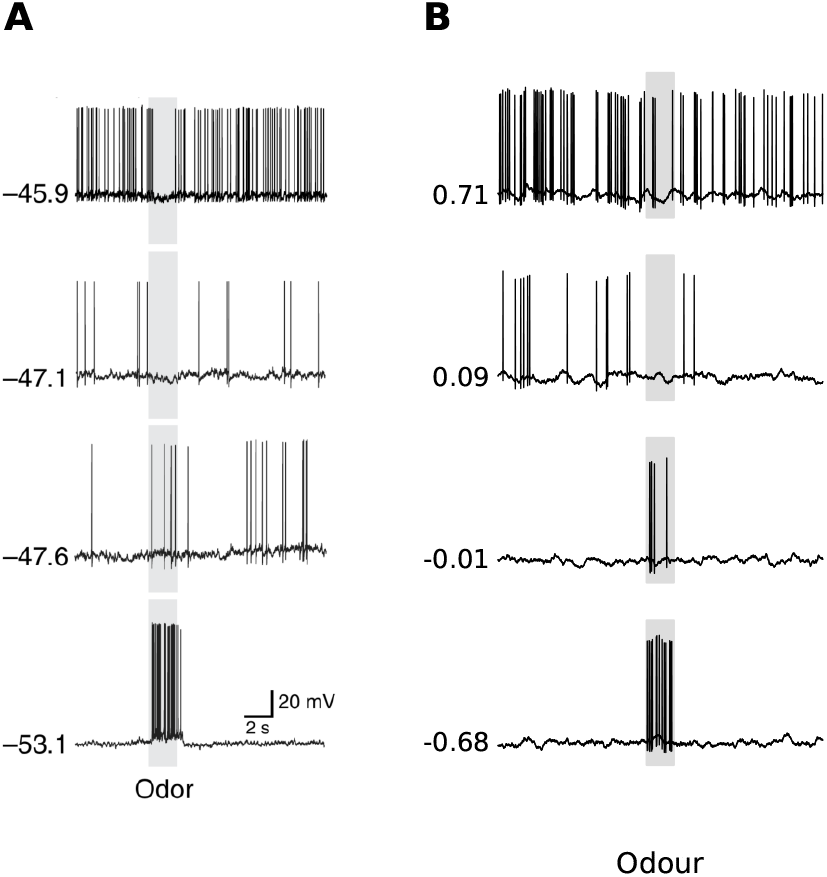
Single cell responses. **(A)** Experimentally observed responses of four cells, ordered by their mean baseline voltages (indicated to the left of each trace). More spikes in the baseline meant fewer spikes in the odour response. Adapted from Fig. 2a of [KSA^+^14]. **(B)** Spiking responses of four simulated cells, ordered by mean baseline voltages (indicated to the left of each trace). Spikes were generated by converting mitral cell voltages to firing rates using a fixed scaling. Noise was added to the resulting traces for display only; it plays no role in the model. More spikes in the baseline meant fewer spikes in the odour response.

Finally, in Figure 2a [KSA^+^14] the authors show the odour evoked responses of a few individual cells, ordered by their baseline activations. The cells with higher baseline activity showed lower odour responses and vice versa. Our mitral cell model uses voltages instead of firing rates, so we could not compare our results directly to these experimental findings. Instead, to perform a speculative comparison, we first converted our mitral cell voltages to firing rates using a fixed scaling factor, and then added noise to the resulting spiking traces to match the levels seen in the experimental findings. In Fig. 8B we plot the resulting ‘spiking’ responses of four cells, selected for their similarity to the experimental results, and ordered by their baseline voltages. The results show good qualitative agreement with the cells shown in [KSA^+^14], suggesting that our model is able to produce spiking responses similar to those experimentally observed.

## 3 Discussion

In this work we have provided a normative explanation for the results of [KSA^+^14] by showing how ‘silent’ mitral cells can emerge as the result of olfactory common feature subtraction.

This computation manifests as the adaptation of the baseline activation of mitral cells to cancel the effect of the common odour or odour features. This in turn requires the baseline activity to reflect the *negative* of the recent activation of each cell, which then explains the negative correlations between baseline and odour-evoked responses found by [KSA^+^14], and reproduced in our simulation results.

### Common feature subtraction as odour-triggered adaptation

Olfactory adaptation is well known from everyday experience and is already present at the receptor level [GS78]. Such adaptation would suffice to remove any *constant* activation caused by background odours. Returning to the motivating example of the wine discrimination, receptor adaptation would remove the effects of the ambient odours of the room. In contrast, CFS is odour-triggered, and only removes components that are common to recently sampled odours. Before receptor adaptation is complete, the background odours in the room would also qualify for common feature subtraction. However, once these odours are removed by receptor adaptation, the only remaining odours available to trigger CFS would be the periodic samples of wine odour, and it would be the common features in these whose negative would ultimately be computed by CFS and stored in the mitral cell baselines. Experience-dependent weakening of mitral cell odour responses has also been observed in awake mice [KCIK12], though over much longer timescales than those considered here.

### Neural implementations

How could the baseline adaptation required by Eq. (17b) be implemented? That equation requires that ***µ***_*i*_ be inhibited by odour-triggered activation of mitral cell *λ*_*i*_, and to excite it in return. Because mitral cells are excitatory, it’s unlikely that ***µ***_*i*_ in this form is implemented by a neuron. One possibility is that ***µ***_*i*_ represents e.g. ionic concentrations that promote a mitral cell’s excitability but are consumed upon activation and replenish slowly. A problem with this interpretation is that such dynamics would be continuously active but those in Eq. (17b) need to be odour-gated.

Another possible implementation flips the sign of ***µ*** in Eq. (17b) and Eq. (17a). In that setting ***µ***_*i*_ is excited by mitral/tufted cell *λ*_*i*_ and inhibits it in turn. Such connectivity motifs are common in the olfactory bulb. One possible implementation would see ***µ***_*i*_ as the summed self-inhibition of a mitral cell by the granule cell spines it contacts [ESM05].

However, such granular feedback seems more complicated than necessary, when all that is needed is negative feedback tied to the mitral cell’s recent activation. Such feedback could instead be provided by one of the periglomerular cells that populate each glomerulus. We leave the determination of the exact neural implementation of the adaptation dynamics to future experimental work.

### Relation to the experimental stimulus set

Our simulations use an explicit common feature: every odour is a binary mixture of a fixed common component and one of eleven unique components (four in Figure 2 and Figure 3). The experiments of [KSA^+^14], by contrast, used a small set of monomolecular odours with no shared component by design. We do not think this is a contradiction. Common feature subtraction does not require that a single shared molecule be present; it requires only that the receptor activations evoked by recently sampled odours share a component that is predictable from one bout to the next— in the simplest case, the mean of the recent odour-evoked activations. Any set of odours that activates overlapping subsets of glomeruli, as monomolecular odours do, has such a component, and the mitral cell baselines should come to cancel it, although we expect the effect to be weaker than with an explicit common odour. The explicit common odour in our simulations makes the computation visible.

### Experimental predictions

Our work predicts, first, that mitral cell background activations are not intrinsic properties of the cells but reflect the *negative* of recent odour evoked activations, and second, that these background activations serve to cancel the contribution of common odour features to the receptor activations, and hence should be inversely related to these. These predictions can be tested by modifying the odour set used in [KSA^+^14] to consist of binary mixtures of different target odours and the same fixed common odour. Repeated exposure of the bulb to a random sequence of these odours according to the protocol in [KSA^+^14] should result in baseline activations being inversely related to odour-evoked responses, as already observed. Our predictions would be (a) that baseline activations would be inversely related to the average of recent odour-evoked responses, (b) that the composition of the ‘silent’ subset of the population would depend *only* on the common odour, and not the target odours; and (c) that the background excitation levels of each mitral cell in the population should be inversely related to the excitation of its parent glomerulus by the common odour. We note that the *sign* of the relationship between baseline and odour-evoked activity is not unique to common feature subtraction: any mechanism that subtracts a running average of a cell’s recent activity would produce it. What is specific to odour-triggered subtraction is the *magnitude* of the relationship. Eq. (44) predicts how the slope depends on the fraction of time odour is present and on the length of the air gaps between presentations. Ungated adaptation, in which the same baseline update is applied at all times rather than only during odour, behaves very differently: during air the baseline then decays with time constant 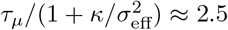 s rather than *τ*_*µ*_ ≈ 20 s, so the spread of baselines largely disappears within a few seconds of clean air and the baseline is only weakly related to the common input (Fig. 4D, grey; fitted slope −0.02, against −0.36 with gating). This yields a directly testable prediction: if subtraction is odour-gated, the spread of mitral cell baselines should persist across odour-free gaps of tens of seconds, decaying over ∼ *τ*_*µ*_.

### Summary

We have proposed a normative explanation for the emergence of ‘silent’ mitral cells in the olfactory bulb, in which these responses emerge as the result of odour-triggered common feature subtraction. Our model gives a circuit-level formulation of olfactory background subtraction consistent with the anatomy of the bulb. Adaptive bulb-like networks that separate the components of fluctuating mixtures have been proposed before [Hop91]; our work differs in deriving the adaptation from an inference problem and in tying it to the baseline activity of individual cells. The model is neurally plausible, is in qualitative agreement with the experimental findings, and can be falsified through a modification of existing experimental protocols.

## 4 Methods

### 4.1 Inference of the unique features in the presence of common features

The probability of the receptor activations in Eq. (13) can be expressed in terms of the activations caused by the common features and, using our Gaussianity assumptions in Eq. (15) and Eq. (16), evaluated analytically as

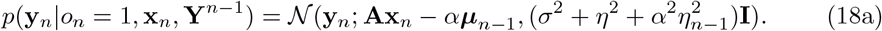

This likelihood is a multivariate Normal distribution just as in Eq. (2). To form the posterior Eq. (12) we combine this term with the prior on **x**_*n*_. We use the same prior as Eq. (1). This means our posterior will have the same form as that in [TSL22], and we can perform the maximization using the dynamics in Eq. (4) but with Eq. (4a) modified to

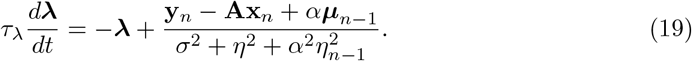

We observe that ***µ***_*n−*1_ now determines the baseline of ***λ***, that is, its activity in the absence of any receptor input **y** or granule cell input **x**.

#### 4.1.1 Updating the effect of the common features

Next, we must use the current observation **y**_*n*_ to update our knowledge of the effects **b**_*n*_ of the common features on receptor activations. We do so in two steps. First, we convert our knowledge in the previous bout, *p*(**b**_*n−*1_ | **Y**^*n−*1^), using the history of observations up to that point, to a prior for the current bout before any observations are made

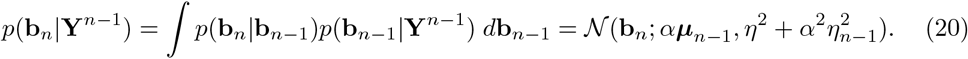

The increased variance reflects the additional uncertainty incurred by the passage of time without a new observation.

Next, we update our prior with the new observation **y**_*n*_ to form a posterior on **b**_*n*_. If we happen to be in an air bout and no odour is present, the new observation does not provide any information about the common features, and the posterior reverts to the prior:

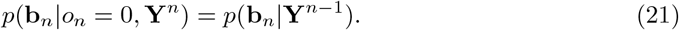

On the other hand, when odour is present, Bayes’ rule weights each **b**_*n*_ by its likelihood to have produced the observation **y**_*n*_,

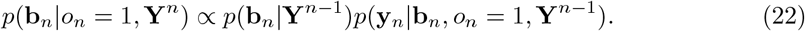

The observations **y**_*n*_ are not produced by the common features **b**_*n*_ alone, but also by the unique features **x**_*n*_. Since the particular unique feature vector responsible for **y**_*n*_ is not given, all possible **x**_*n*_ must be accounted for, weighted by their priors

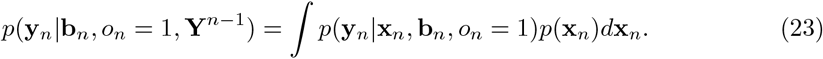

The form of the prior term, the same as Eq. (1), makes computing the integral in Eq. (23) difficult. To approximate it, we use the fact that the granule cell dynamics of Eq. (4b) and Eq. (4c), driven by the mitral cell dynamics in Eq. (19) yield the MAP estimate 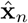 of the unique features for the case that odour is present. So, rather than considering every possible **x**_*n*_ to compute the likelihood in Eq. (23), we consider only the MAP estimate

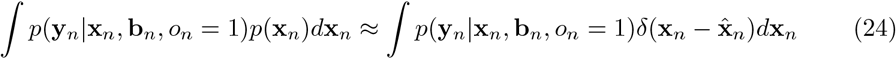

to arrive at

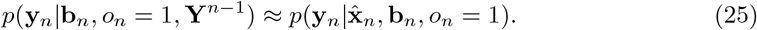

Substituting this and Eq. (20) into Eq. (22) yields

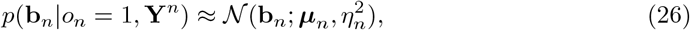

where the updated mean and variance are

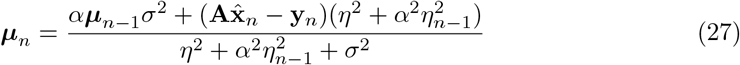

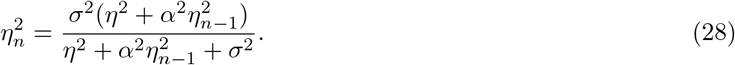

### Computing the mean update

The mean update of Eq. (27) can be expressed in terms of the mitral cell activations

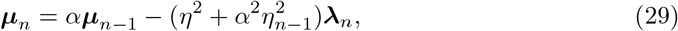

where ***λ***_*n*_ is what the mitral cell dynamics of Eq. (19) converge to at the end of the bout. Collecting Eq. (20), Eq. (21) and Eq. (29) we get

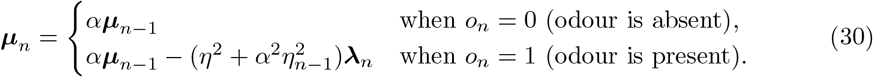

which we can combine into a single update rule

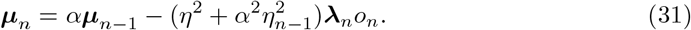

Notice that the contribution of the mitral cell activations ***λ***_*n*_ is gated by presence of odour as indicated by *o*_*n*_.

### The variance update

Similar to Eq. (30), we collect Eq. (20), Eq. (21) and Eq. (28) as

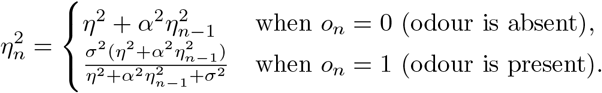

and express this as a single update

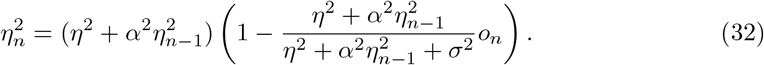

Rather than model this variance update directly, we approximate by the steady-state value 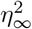 achieved when we replace *o*_*n*_ by its expectation *ō*. Replacing 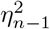 and 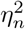 with 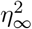 in Eq. (32), we get

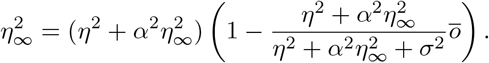

To solve this, we define 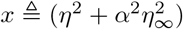. Multiplying both sides of the equation above by *α*^2^, we get

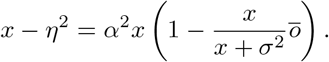

Solving for *x*, we get

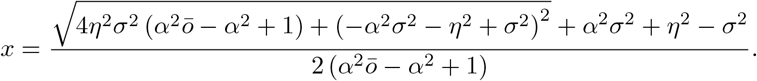

Converting from *x* back to 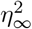, we get

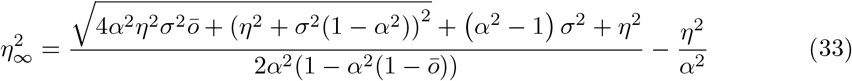

### 4.2 Mapping discrete-time dynamics to continuous-time

To map the discrete-time update in Eq. (31) to continuous time, we view them as the activity of a first order system subjected to step input at the beginning of the bout (*t* = 0)

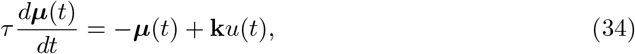

where *u*(*t*) is the unit step function, reported at the end of the bout (*t* = *T* ):

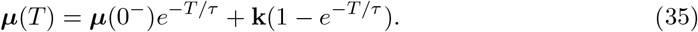

Setting ***µ***(*T* ) to ***µ***_*n*_ and ***µ***(0^*−*^) to ***µ***_*n−*1_ we see that

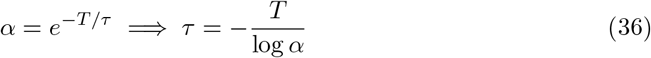

and

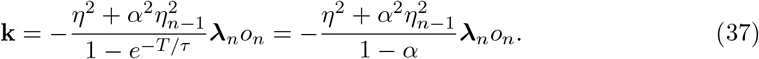

Replacing ***λ***_*n*_ and *o*_*n*_ by their continuous time values we arrive at Eq. (17b); with 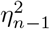 replaced by its steady-state value 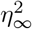 and a factor of *α* absorbed into the redefinition of ***µ***, this step amplitude is **k** = −*κ* ***λ***_*n*_*o*_*n*_, the origin of the *κ* in Eq. (17b).

### 4.3 Simulation details

In the simulations the *M* ×*N* mixing matrix **A** has independent standard-normal entries, each column normalised to unit norm. Each target odour and the common feature is a single molecular species (a one-hot vector) placed on an evenly spaced grid, the common feature scaled to concentration 0.9. Time is divided into one-second bouts: an odour bout presents a binary mixture of the common feature and one target odour (targets cycled in turn), and successive odour bouts are separated by 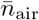 air bouts. We integrate the system dynamics (Eq. (17b) and accompanying equations) with an Euler step Δ*t*, holding the innovation variance at its steady-state value 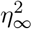 (Eq. (33)). The odour response of a cell is its mean voltage over the one-second presentation and its baseline response is its mean voltage over the preceding 500 ms. All parameters of the default run—used for Figures 4–7—are listed in Table 1; Figures 2 and 3 use the same parameters with a reduced set of four target odours. Figure 8 uses a separate run with the same parameters except 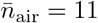; the four cells shown there were selected for their resemblance to the cells in Fig. 2a of [KSA^+^14]. The baseline update (Eq. (17b)) is switched on only after one priming presentation of each target odour, so the early and late states in Figure 2 are before and after this point. The receptor noise *σ* and innovation noise *η* enter the simulations only as parameters of the generative model, through 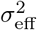 and *κ*: the simulated receptor input is noiseless during odour bouts and carries only small Gaussian noise (s.d. ≈0.014 per glomerulus) during air bouts, and the simulated common feature is constant across bouts. In all simulations *o*(*t*) is supplied directly rather than inferred. The repository at https://github.com/stootoon/silent-mcs-release contains code to reproduce all figures.

### 4.4 Slope of mitral cell baseline against common input

We’re interested in the relationship between the mitral cell baseline, ***µ*** and the common input it receives, **Ac**. To determine this, we start with our fixed-point equation for the mitral cell activity, where **x**^*∗*^ denotes the true unique-feature vector of the presented odour,

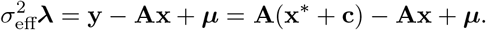

Averaging over odours, we get

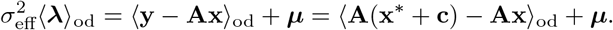

Notice that the baseline retains its value over the averages. We assume that our estimate satisfies

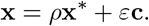

The coefficients *ρ* and *ε* will in general depend on **x**^*∗*^ and **c**. Our average reconstruction error then becomes

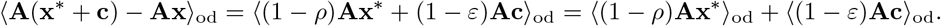

Given a large and diverse enough odour set, we expect the first term to average to 0. For the second term, we assume that *ε* and **c** are uncorrelated over odours and so can be averaged separately. Abusing notation, we’ll use *ε* for its odour average as well, so that our average reconstruction error becomes

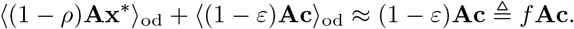

The fixed-point equation for the average mitral cell activity then becomes

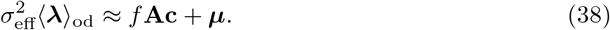

The baseline activity integrates over odour presentations, so

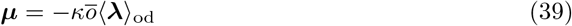

We can substitute Eq. (38) into Eq. (39) to relate ***µ*** to **Ac**,

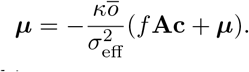

Rearranging, we get the relationship as

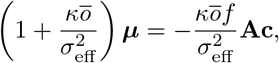

so that

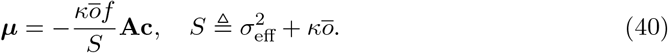

We need a third equation to solve for *f* . To provide that, we consider the GC activity for the common odour. Since the common odour is a single molecular species at concentration *c*, we can write **Ac** = *c* **a**, where **a** is the column of **A** for that species (its receptive field); the activity *x*_*c*_ of the corresponding granule cell then satisfies

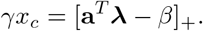

Its odour average is

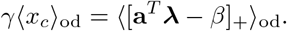

If we assume that the projection is large enough that the rectification can be ignored, we can write this as

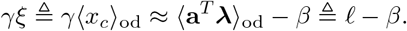

The projection of the baseline satisfies

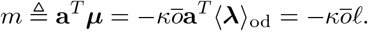

The projection of the mitral cell activity is

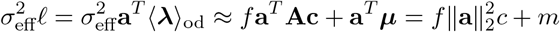

Using

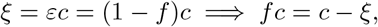

we can list our system of equations as

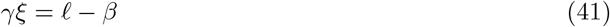

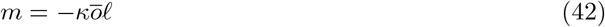

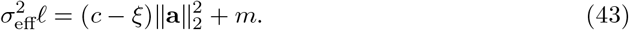

Substituting Eq. (42) into Eq. (43) gives

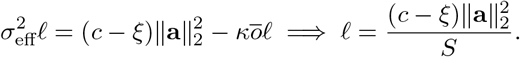

Substituting that into Eq. (41) gives

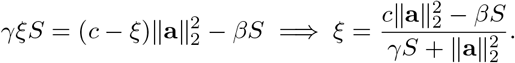

Then,

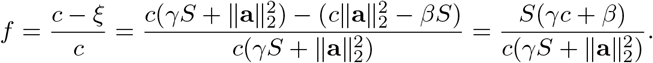

Plugging this back into Eq. (40), we arrive at

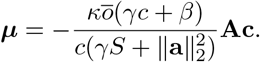

This gives the average relationship during odour presentations. The baseline activity exponentially decays in between presentations. If we take the average duration of an air gap to be 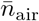 bouts then the decay is 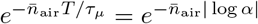, so our relationship between the *pre-odour* baseline and the background input becomes

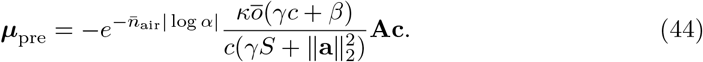

### 4.5 Slope of odour-evoked response against baseline

In air, the glomerular input is approximately zero and the granule cells are below threshold, so by Eq. (17a) the pre-odour mitral cell voltage is 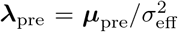. Combining Eq. (39) with the pre-odour decay 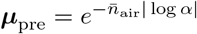 gives

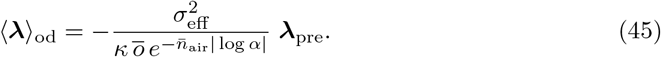

The odour-averaged response of a mitral cell is therefore predicted to be a linear function of its pre-odour voltage, with a slope that depends only on the noise variances, the learning gain, and the odour statistics, and not on the granule cell parameters *β* and *γ*. This is the relationship plotted in Fig. 5C; individual cell-odour pairs scatter about it by the odour-specific residual.

### 4.6 Inferring the presence of odour

Inferring the unique odour in the current bout **x** given the history **Y**^*n*^ of receptor observations requires knowing whether odour is present in the current bout or not, since

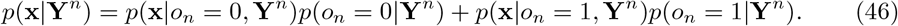

One way to determine the presence of odour from the history of receptor observations is to use the system’s current best estimate of the unique and common odours. The posterior on odour being present is

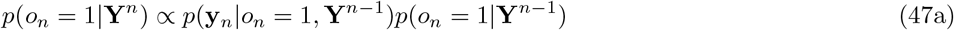

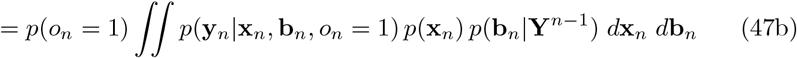

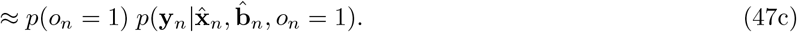

Here we have assumed that the presence of odour in successive bouts is independent, so that *p*(*o*_*n*_ = 1|**Y**^*n−*1^) = *p*(*o*_*n*_ = 1). The posterior on it being absent is

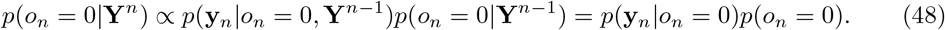

Then the ratio of posteriors is

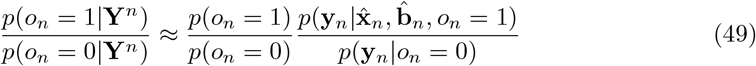

So the relative probabilities of odour vs. air can be approximated from the results of the inference calculation that assumes *o*_*n*_ = 1. Alternatively, we can assume that *p*(*o*_*n*_ = 0|**Y**^*n*^) is some function of some statistic of **y**_*n*_, such as its variance, that is built into the system via e.g. the firing-rate nonlinearity of a neuron. In the simulations reported here *o*(*t*) is supplied directly.

## Acknowledgements

We are grateful to Paul Masset, Jacob Zavatone-Veth, the members of the Schaefer lab at the Crick, and the Latham lab at the UCL Gatsby Unit for feedback on this manuscript. This work was supported by the Francis Crick Institute which receives its core funding from Cancer Research UK (FC001153), the UK Medical Research Council (FC001153), and the Wellcome Trust (FC001153); and by the NSF/CIHR/DFG/FRQ/UKRI-MRC Next Generation Networks for Neuroscience (“NeuroNex”) Program “From Odor to Action” (Award #2014217).

## References

[ESM05] Veronica Egger, Karel Svoboda, and Zachary F. Mainen. Dendrodendritic Synaptic Signals in Olfactory Bulb Granule Cells: Local Spine Boost and Global Low-Threshold Spike. The Journal of Neuroscience, 25(14):3521–3530, June 2005.

[GBB+17] Agnieszka Grabska-Barwinska, Simon Barthelmé, Jeff Beck, Zachary F. Mainen, Alexandre Pouget, and Peter E. Latham. A probabilistic approach to demixing odors. Nature Neuroscience, 20(1):98–106, January 2017.

[GS78] T. V. Getchell and G. M. Shepherd. Adaptive properties of olfactory receptors analysed with odour pulses of varying durations. The Journal of Physiology, 282(1):541–560, 1978.

[Hop91] J. J. Hopfield. Olfactory computation and object perception. Proceedings of the National Academy of Sciences, 88(15):6462–6466, 1991.

[KCIK12] Hiroyuki K. Kato, Monica W. Chu, Jeffry S. Isaacson, and Takaki Komiyama. Dynamic sensory representations in the olfactory bulb: modulation by wakefulness and experience. Neuron, 76(5):962–975, 2012.

[KR11] Alexei A. Koulakov and Dmitry Rinberg. Sparse Incomplete Representations: A Potential Role of Olfactory Granule Cells. Neuron, 72(1):124–136, October 2011.

[KSA+14] Mihaly Kollo, Anja Schmaltz, Mostafa Abdelhamid, Izumi Fukunaga, and Andreas T. Schaefer. ‘Silent’ mitral cells dominate odor responses in the olfactory bulb of awake mice. Nature Neuroscience, 17(10):1313–1315, October 2014.

[PBAP14] Il Memming Park, Yuriy V. Bobkov, Barry W. Ache, and José C. Príncipe. Intermittency coding in the primary olfactory system: a neural substrate for olfactory scene analysis. Journal of Neuroscience, 34(3):941–952, 2014.

[RHKM14] Dan Rokni, Vivian Hemmelder, Vikrant Kapoor, and Venkatesh N. Murthy. An olfactory cocktail party: figure-ground segregation of odorants in rodents. Nature Neuroscience, 17(9):1225–1232, 2014.

[TSL22] Sina Tootoonian, Andreas T. Schaefer, and Peter E. Latham. Sparse connectivity for MAP inference in linear models using sister mitral cells. PLOS Computational Biology, 18(1):e1009808, January 2022.

[ZVMT+23] Jacob A. Zavatone-Veth, Paul Masset, William L. Tong, Joseph D. Zak, Venkatesh N. Murthy, and Cengiz Pehlevan. Neural circuits for fast Poisson compressed sensing in the olfactory bulb. In Advances in Neural Information Processing Systems, volume 36, pages 64793–64828, 2023.

